# Screening of beans (*Phaseolus vulgaris* L.) genotypes for drought tolerance

**DOI:** 10.1101/2021.01.14.426641

**Authors:** Jigme Thinley, Choki Dorji

## Abstract

Drought is an important factor limiting crop yield and food insecurity globally. Sixty- percent of the bean production areas are prone to drought and subsequently result in eighty-percent of yield reduction. It is emerging as a rising threat to farming communities in Bhutan. Limited studies on crop drought tolerance done in Bhutan. Six bean genotypes (Orey serbu, Orey regtang, Orey brokchilu, Yadhipa orey, Kerongree orey, Brokopali) chose to ascertain drought tolerance. The study was a factorial experiment with Completely Randomized Design (CRD) with six treatments and three replications. Genotypes were subjected to drought stress after 50% flowering till the start of pod formation. The drought condition induced for ten days during the pod formation and the irrigation resumed till harvesting. Means of water-stressed genotypes compared to their corresponding non- stressed with an All-pairwise comparison using the Bonferroni test at the significant level p<.05. The growth (leaf area, relative water content, root length, shoot weight, maturity) and yield parameters (the number of pods, pod length, the number of seeds number per pod, seed weight) determined during the time of harvest. There were significant differences (p<.05) in all the parameters measured under stress and non-stress conditions. Water-stress decreases plant growth and development of all the bean genotypes. With regards to drought susceptibility index (DSI), Orey serbu(1), Orey regtang(1), Yadhipa orey(1), and Kerongree orey(1) had the lowest DSI value. The lesser the susceptible indices, the greater is the tolerance level to the drought and vice versa.

## 1. Introduction

Common beans (*Phaseolus vulgaris* L.) consume as a staple and low-cost protein source in the underdeveloped countries where protein malnutrition is prevalent [13]. [12] stated that the genus *Phaseolus* originated from the Mesoamerican regions comprises five domesticated species: *Phaseolus vulgaris* (common bean), *Phaseolus dumosus, Phaseolus coccineus* (runner beans), *Phaseolus acutifolius* (tepary beans), and *Phaseolus lunatus* (Lima bean). Dry beans are a staple food in Latin American, Eastern, and South African countries [2]. Similarly, beans are the source of dietary carbohydrates and micronutrients for more than 300 million people from Eastern Africa and Latin America [50, 9]. India produced 6,75,188 metric tons [MT] of green beans, followed by Bangladesh (137495 MT) in 2017 [21**Error! Reference source not found.**]. As per [16], Bhutan produced 1,475 MT of beans in 2016 and 5,273 MT in 2017 [17]. Tsirang has the highest harvested area (475 acres), and Bumthang the least harvested area (3 acres) [17]. Bean production is adversely affected by both biotic and abiotic factors [23], and the major abiotic factor affecting crop production worldwide is water stress [28]. Water-stress is ranked second after pests and diseases in reducing the agricultural grains. Sixty-percent of the bean production areas are prone to the drought that drains an eighty-percent of yield [24, 41]. The 60% of beans production in the world occurs in the agricultural land prone to water deficit, lack of irrigation system, and the dry periods may result in an 80% yield reduction [41]. According to [17], the drought (4% of total households) and the insufficient irrigation supply (27% of total households) are emerging as a threat to agriculture in Bhutan. Beans are the most important, directly consumed food legume, and it is one of the most important cash crops after potato grown by every Bhutanese farmer. Almost every Bhutanese household consumes beans and remains as highly demanded in the market; however, the study by [49] reported that the drought gets severer in the future due to continuous anthropogenic activities. This study aimed to identify the drought-tolerant bean genotypes, which would help maintain the production and to meet the demand in the Bhutanese market, and provide the basis for the development of drought-tolerant hybrids in the future.

## 2. Materials and Methods

### 2.1 Study site, experimental design, stress treatment, and irrigation schedule

The greenhouse of the agriculture farm of College of Natural Resources (CNR), Lobesa, was used to perform this study in August 2019. It is at an elevation of 1450 meters above sea level between 27° 30’ 1’’ N and 89° 52’ 42’’ E of Greenwich. The area experiences an annual rainfall of 500 mm to 1500 mm with temperatures ranging from 5℃ to 30℃. It lies in the dry sub-tropical region and experiences hot and humid summers during the monsoon months of June, July, and August. The bean varieties used for the experiment were Yadhipa orey(13), Orey serbu(27), Kerongree orey(7), Orey regtang(6), Orey brokchilu(15), and Bropali(25). Used completely randomized design (CRD) for this factorial experiment. There were six treatments (six bean genotypes) and three replications (three plants per replica) and classed treatments into the water-stressed and nonstressed groups. Used a total of 108 potted plants during the experiment, and the distance between the pots was 30 cm within rows and 40 cm between rows [4]. For media, used 3 kg of topsoil to fill 4,416 cm3 volume plastic pots. Subjected genotypes to drought stress after 50% flowering and lasted for ten days, and resumed irrigation till harvesting similar to [4, 8]. Weighed pots following one-day intervals [15] and applied the water by calculating the required amount of water using the formula; the amount of water = the weight of pot at field capacity - pot weight at field condition and maintained the soil moisture near field capacity (FC) throughout the experiment [8].

### 2.2 Data collection and analyses

Recorded the weather data using Datalogger (temperature and relative humidity) and recorded both growth and yield attribute data during physiological maturity that defines the number of days for 75% to 90% of pods to lose their green pigmentation [48]. Collected data on leaf area, the number of pod per plant, weight of seed per plant, shoot dry weight, root dry weight, and calculated drought susceptibility index (DSI).

Calculated drought Susceptibility index (DSI) using the following formula:

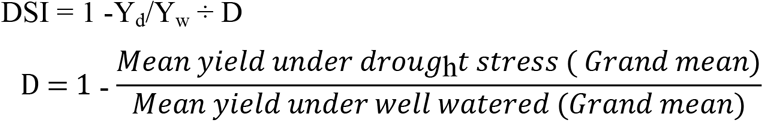

Where

D = drought intensity

Y_d_ = Mean yield under drought condition (Average yield of genotypes

Y_w_ = Mean yield under well-watered condition. (Average yield of each genotype)

Data were subjected to a two-way analysis of variance (Two-way ANOVA) to compare differences in the means. Calculated statistical significances at p < .05 and collated group means following Bonferroni pairwise comparison tests. The factorial design in statistical program Statistix version 8 used for all the analyses.

## 3. Results and Discussion

### 3.1 Effect of drought stress on leaf area (dm^2^)

The significant difference in leaf area was recorded between the treatments group (stress and non stress) [*F* (1, 22) = 52.96, *p* < .05] and within the genotypes group [*F* (5, 22) = 10.42, *P < .*05]. There was a significant interaction between the treatment and genotypes (*p* < .05). Results indicate the genotype difference in leaf area was affected by the drought stress. Under the water-stressed condition, most of the genotypes had a smaller leaf area than non-stress [Figure 1]. Orey serbu (*M* = 0.55 dm^2^, *SD* ± 0.05) and Orey regtang (*M* =0.55 dm^2^, *SD* ± 0.03) recorded significantly higher in leaf area under water stress. However, there was no significant difference (*p* > .05) between genotypes Yadhipa orey, Orey brokchilu, and brokpali in stress conditions. Wilting, shedding, and curling of leaves, closure of stomata, and reduction of cell enlargement could be the reason for the reduction in leaf area under the stress condition. During the onset of water stress, it inhibits cell elongation in the leaf, and the lower leaf area leads to less water uptake from the soil, and transpiration is reduced [20]. Under the non-water stressed condition genotypes, Orey serbu had recorded significantly higher in leaf area (*M* = 0.75 dm^2^, *SD* ± 0.01). The result coincided with [42, 15], who observed that plants grown under water deficit conditions showed a lower leaf area than plants grown under control conditions. [43] argued that the leaf area is related to plant metabolism, dry matter production, and yield.

**Figure 1:**
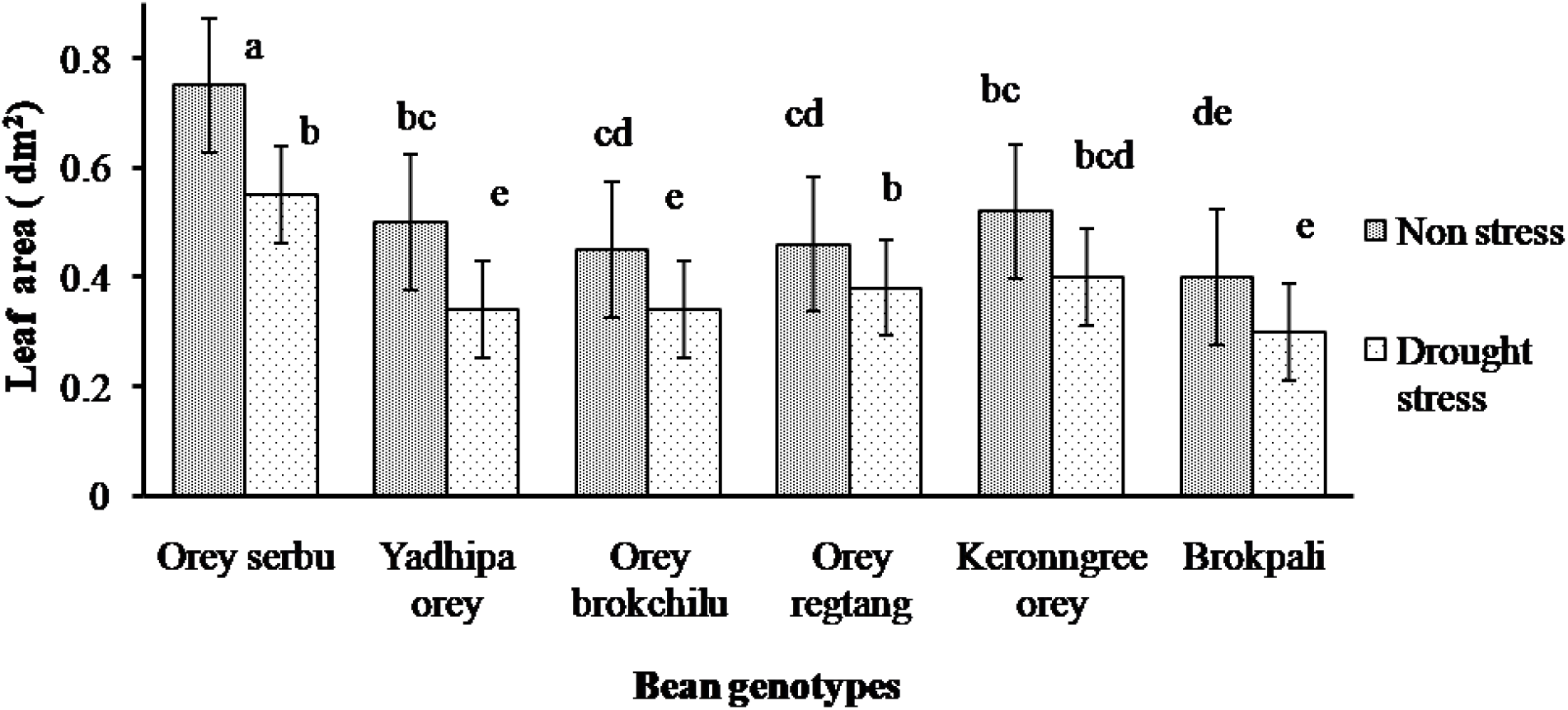
Leaf Area (dm^2^).

### 3.2 Effect of drought stress on pod number

There was a significant difference in the number of pods per plant in both between treatments and genotypes group [*F* (1, 22) = 64.61, *P* < .05] [*F* (5, 22) = 8.67, *p* < .05]. The study revealed the significant (*p* <.05) interaction between treatment and genotypes and shows that the varietal difference in pod number was affected by stress imposed [Figure 2]. Under the water-stressed condition, the number of pods per plant decreased compared to non-stress. The highest average number of pods was in Yadhipa orey (*M* = 5.25, *SD* ± 0.33). The fewer floral part formation could have affected the number of pods per plant. Under the non-water stressed condition, Orey regtang had the highest (*M* = 7.39, *SD* ± 0.98) average number of pods per plant. The present findings are in line with [13], who reported that under the high moisture stress during the reproductive stage, the plants are exposed to floral abortion resulting in low yield. Other authors [7, 44] also reported that water-stress imposed during the flowering and pod setting causes flower and pod abortion. The result agrees with the finding of [7, 42, 5] and [29] reported the decline in pod number when irrigated in 21 days intervals. According to [33], the reproductive stage is the most sensitive to drought stress.

**Figure 2:**
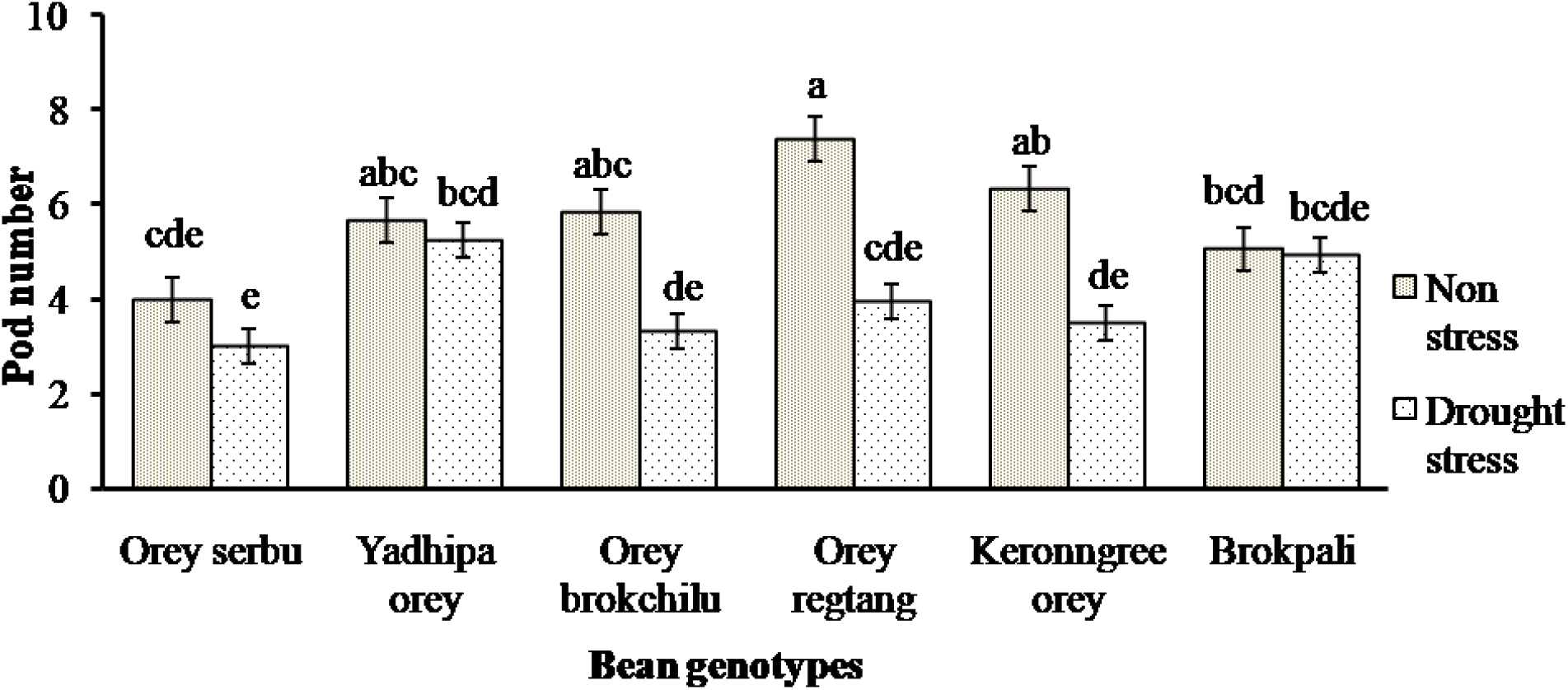
Number of pods per plant.

### 3.3 Effect of drought stress on seed weight (g)

There was a highly significant difference in seed weight both between treatment and within genotypes group [*F* (1, 22) = 597.64, *p* < .05] [*F* (5, 22) = 35.76, *p* < .05]. Under the stress condition, seed weight/yield decrease compared to non-stress [Figure 3]. However, Brokpali recorded the highest seed weight (*M* = 5.94g, *SD* ± 0.59). The reduction in seed weight/yield under the water stress was may be due to a decrease in carbohydrate assimilation that might have lead to less pod number. Under the non-water stress condition, Brokpali had the highest average seed weight of (*M*= 14.51g, *SD* ± 1.51). The reduction in a legumes seed yield under the stress condition was due to the lower number of pods per plant [27, 38, 30, 7]. The reduction in the seed yield and the number of pods is due to the detrimental effects of drought on pod set and grain filling [14].

**Figure 3:**
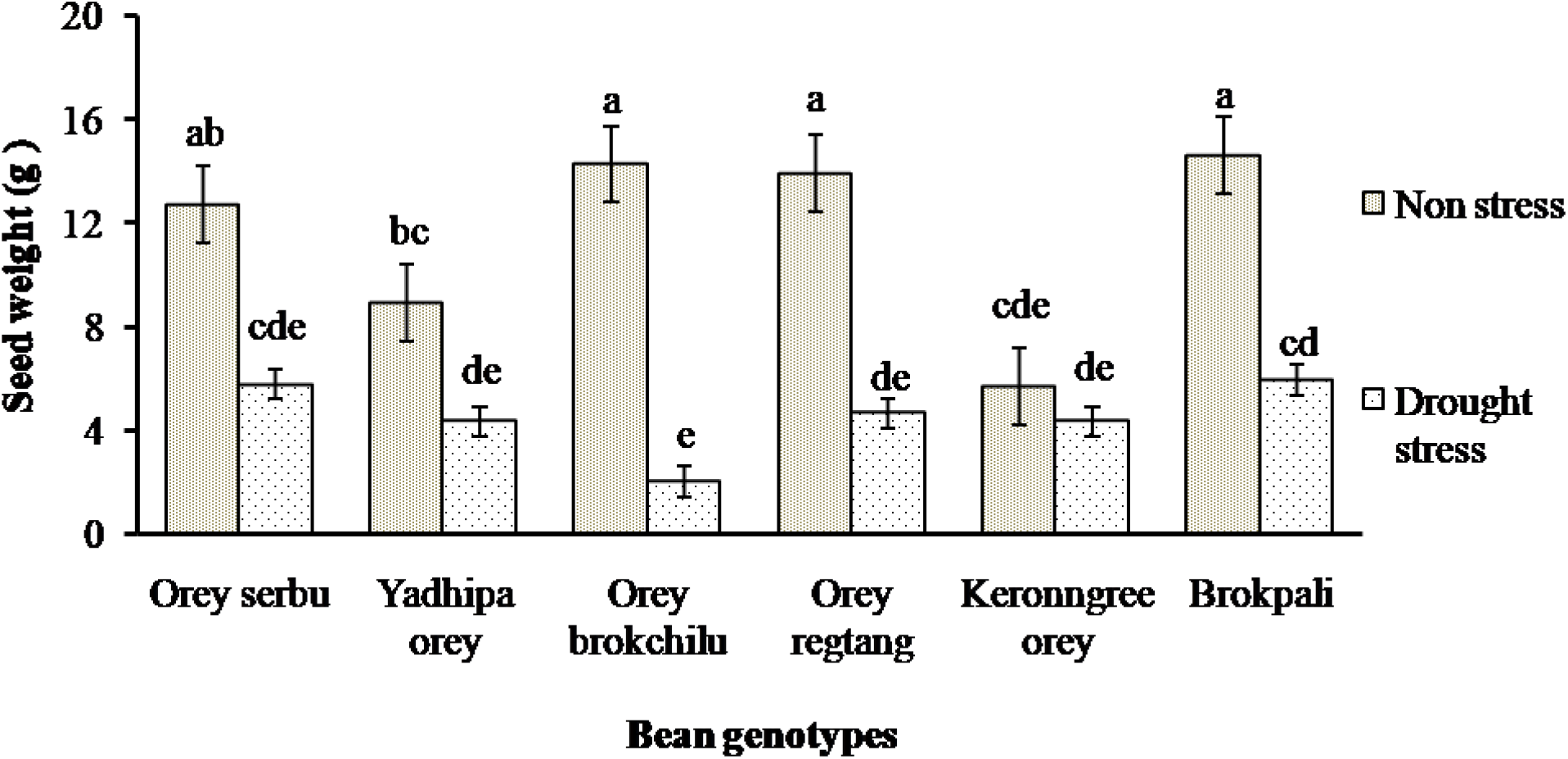
Seed weight (g) per plant.

A similar result obtained by [46], yield reduction in common beans of 58-87% occurs when the water stress is set forth during its reproductive stage. The result accords with the findings of [32, 45, 3, 1]. [24] also had a similar observation in physiological responses of common bean (P*haseolus vulgaris* l.) genotypes to water stress. Moreover, reported seed yield reduction under water stress for a black gram and green gram [47], soybean [26], and Phaseolus vulgaris [36, 22, 15]. The lower seed yield in common beans under stress is due to the lower photosynthate assimilation and carbohydrate partitioning to the developing grain due to drought stress [37, 40, 6].

### 3.4 Effect of drought stress on dry weight (g) of the shoot

The study revealed a significant difference in shoot weight between the treatment group [*F* (1, 22) = 40.15, *p* <.05] and the genotypes group [*F* (5, 22) = 13.81, *p* <.05]. Under the water stress condition, plant shoot dry weight decreased significantly than non-stress [Figure 4]. It could be due to limited photosynthate assimilation and the limited amount of water absorbed by roots. Orey serbu was observed the highest shoot weight of (*M* = 6.91 g, *SD* ± 0.14) under water stress.

**Figure 4:**
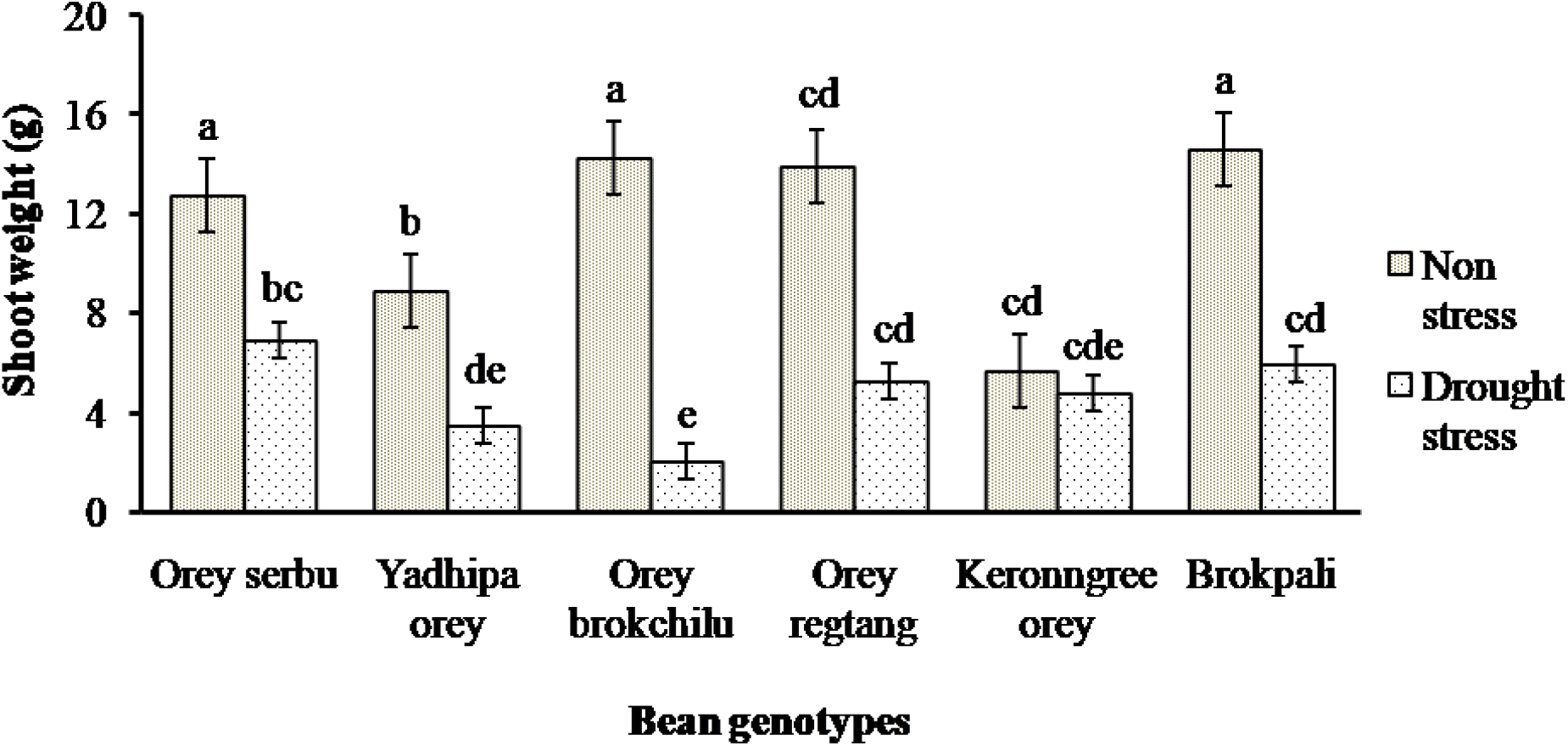
Shoot weight (g)

For non-water stress conditions, Brokpali had the highest shoot weight (*M*= 14.61 g, *SD* ± 0.29). A similar finding was investigated by [35, 10] there is more reduction in shoot growth than root growth under stress. According to [46], both shoots and roots are the most affected parts of plants and key components of the adaptation to drought at the morphological level.

Drought stress severely reduces stem growth and the ability to intercept solar radiation [1]. Among the genotypes, in stress conditions, Orey serbu, Brokpali, and orey regtang showed significantly higher shoot dry weight than the other genotypes. According to [11], a positive correlation (r = 0.53**) exists between shoot and root dry weight in optimal irrigation. However, no correlation (r =0.11) exists under drought stress. It could be due to the genotypic difference in terms of root adjustment to stress.

### 3.5 Effect of drought stress on root dry weight (g)

Significant difference (*p* <.05) was observed in the root weight both between the treatment and genotypes group [*F* (1, 22) = 135.47, *p* < .05] [*F* (5, 22) = 6.36, *p* < .05]. Under water-stressed conditions, root dry weight decreased compared to non-stress [Figure 5]; however, Brokpali had the highest root weight (*M* = 1.84 g, *SD* ± 0.36) under water stress. The decreased in root dry weight could be due to poor roots proliferation in a stressful environment. Plants grown under water deficit conditions had low root biomass [34]. For non-water stress conditions, Keronngree orey recorded the highest root weight (*M* = 2.9 g, *SD* ± 0.23).

**Figure 5:**
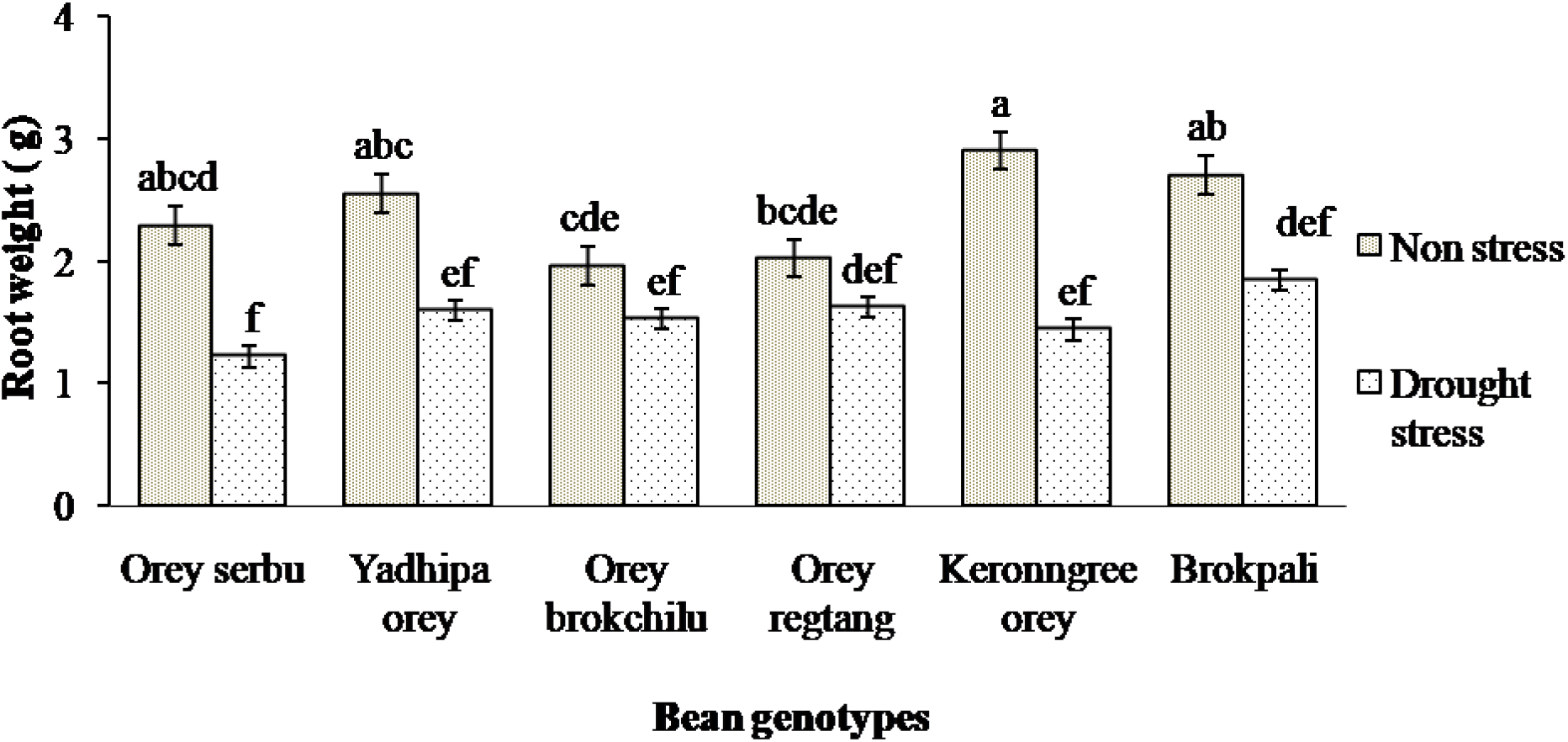
Root weight (g).

Yield components and Biomass accumulation in beans are affected by drought stress [31]; however, few genotypes under the stress condition seem to perform better (Brokpali (1.84 g), Orey regtang (1.62 g), and Yadhipa orey (1.60 g)). It could mean that these genotypes have better soil water acquiring efficiency compared to other genotypes. According to [18], the number of roots decreases with a decrease in water potential. Moreover, reduction in lateral roots directly affects the root biomass [39]. The root characters (biomass, length, density, and depth) are the main drought avoidance traits that contribute to the final yield under terminal drought environments [25].

### 3.6 Drought intensity (D) and Drought susceptibility Index (DSI)

Determined the drought susceptibility index (DSI) on relative water content, shoot weight, seed weight, root weight, leaf area, pod number, pod length, seed number, and root length for six water-stressed genotypes. The Drought susceptibility index (DSI) for [Table 1] leaf area ranged from −0.27 to 0.27, −0.13 to 1.45 for shoot weight, 0.56 to 1.43 for root weight, 0.05 to 2.14 for relative water content, 0.38 to1.41 for seed weight, −0.41 to 0.22 for pod number, −0.06 to 0.05 for pod length, −0.23 to 0.2 for seed number, and −0.32 to 0.17 for root length. The smaller the value of the drought susceptibility index, the more tolerant is the genotypes to water stress conditions.

**Table 1:**
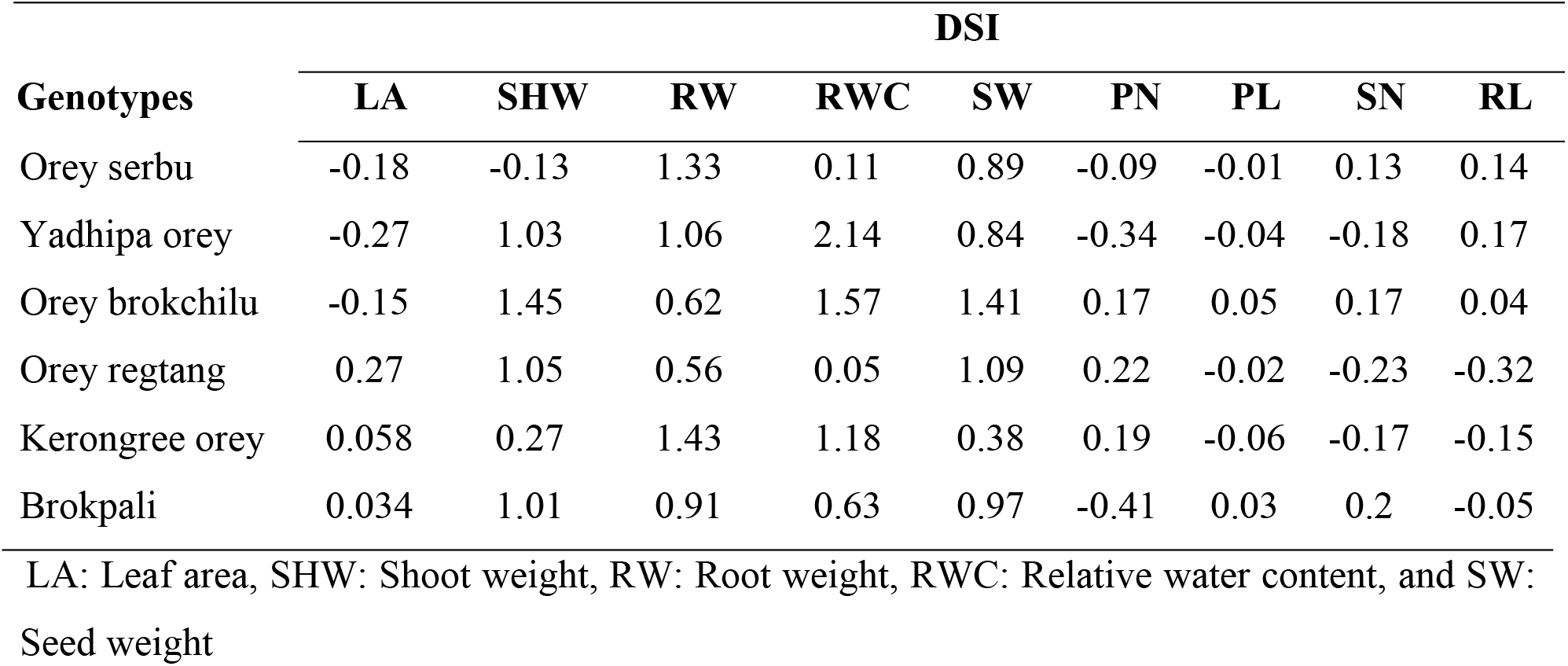
Drought susceptibility index (DSI)

### 3.7 Scoring and ranking of genotypes based on drought susceptibility indices

Ranked the water-stress toleration of six bean genotypes [Table 2]; the scoring and ranking were such that the least susceptible genotypes scored 1, and the highest susceptible genotypes score 6 [Table 4]. Values of Drought Susceptibility Index (DSI) less than 1.0 indicate toleration to drought. Values of DSI equal to 0.0 indicate maximum possible drought tolerance (no effect of drought on yield). Four genotypes (Orey serbu, Yadhipa orey, Orey regtang, and Kerongree orey) had the least DSI (rank 1st) and have a greater drought tolerance level. Orey brokchilu had the highest DSI (ranked 6th), followed by Brokpali (ranked 5th) and more susceptible to drought conditions.

**Table 2:**
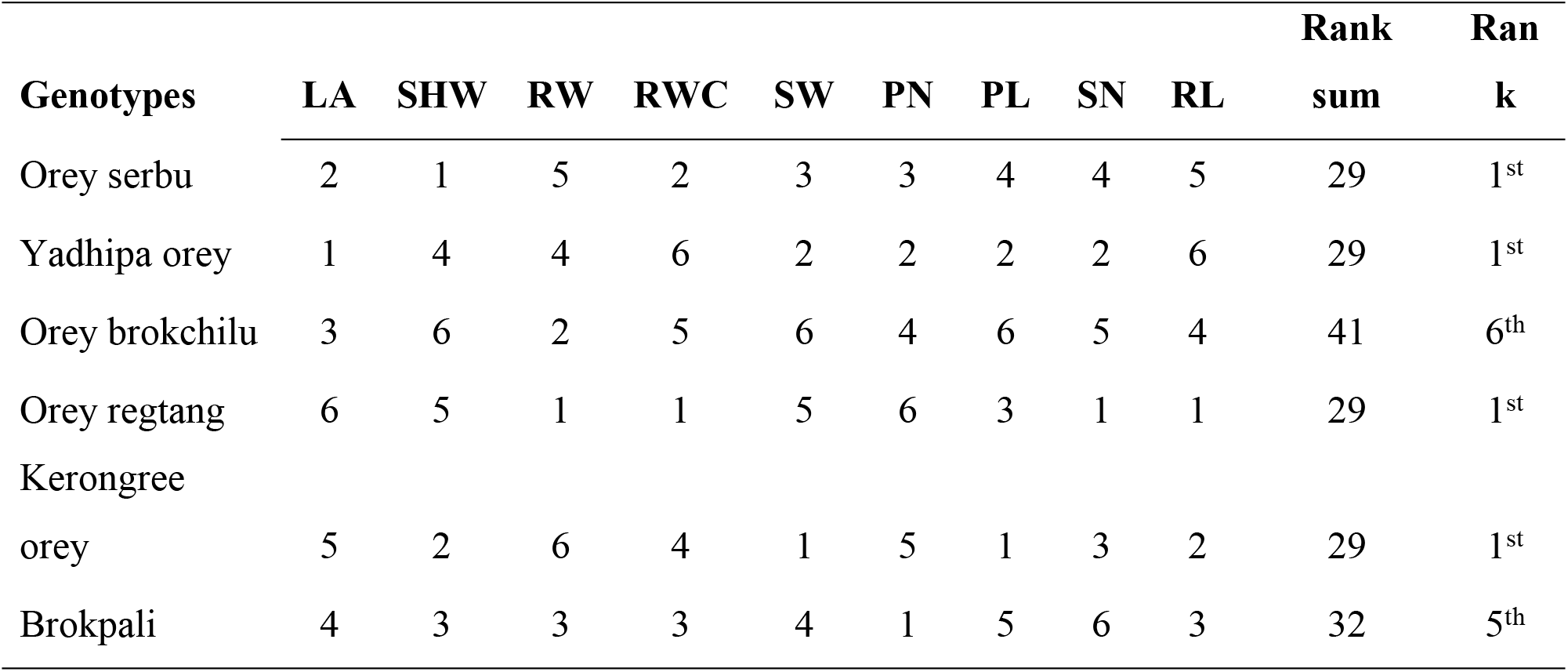
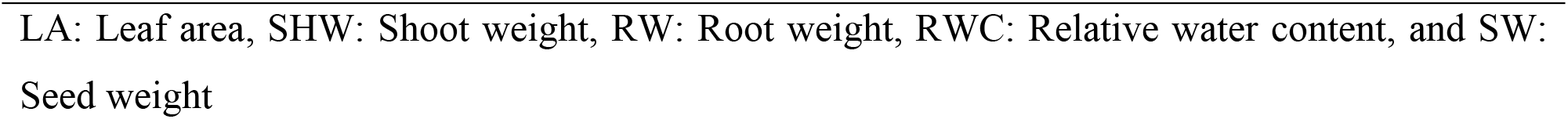
Scoring and ranking of genotypes.

## Conclusion

The present study revealed Orey serbu (rank 1st), Yadhipa orey (rank 1st), Orey regtang (rank 1st), and Kerongree orey (rank 1st) would perform better under conditions of limited water supply based on their low drought susceptibility index in the greenhouse. Brokpali (rank 5th) and Orey brokchilu (rank 6th) were least tolerant to drought. Leaf area, the number of pods, the number of seeds, and seed weight per plant was significantly reduced by drought stress compared to non-stress. Drought stress affected the proper growth and development and ultimately affected the yield of the plants.

## Acknowledgment

We would like to extend our heartfelt gratitude to the Department of Agriculture, College of Natural Resources, and lab assistant for their supervision, suggestion, advice, guidance, encouragement, and commitment during the research work. Lastly, we acknowledge lecturers, friends, and my family for supporting our study.

## Supporting Documents

**Table 3:**
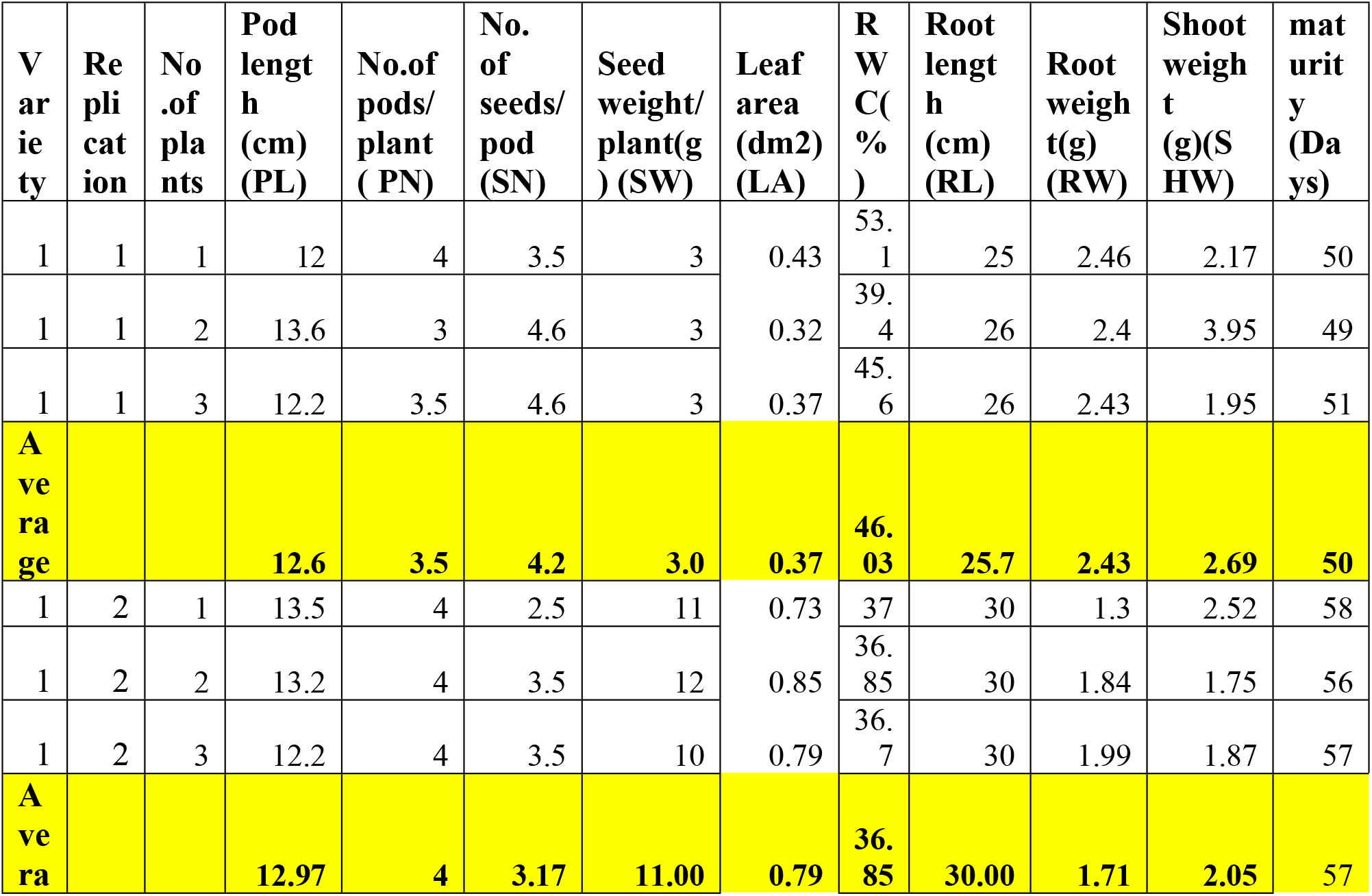
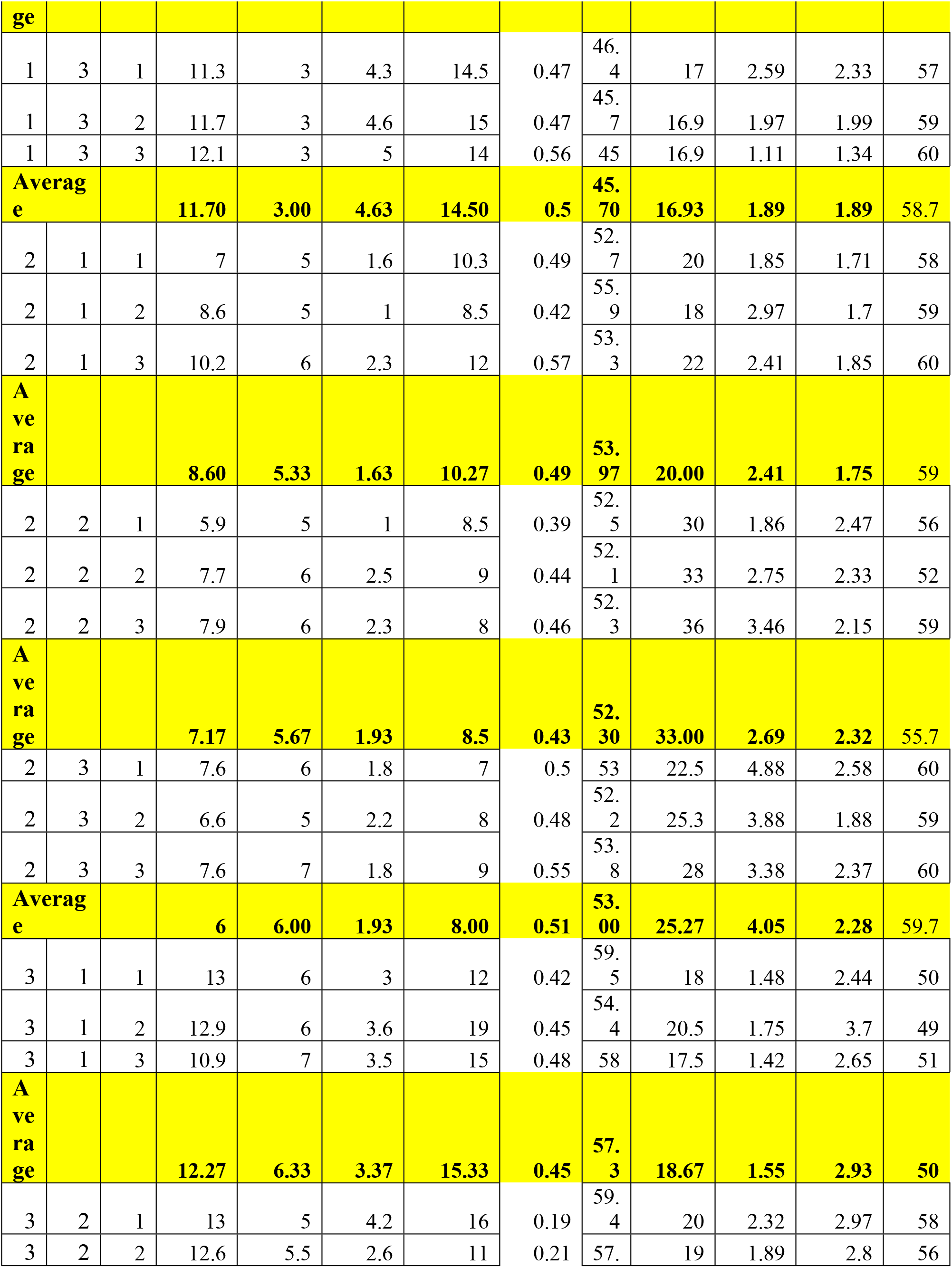
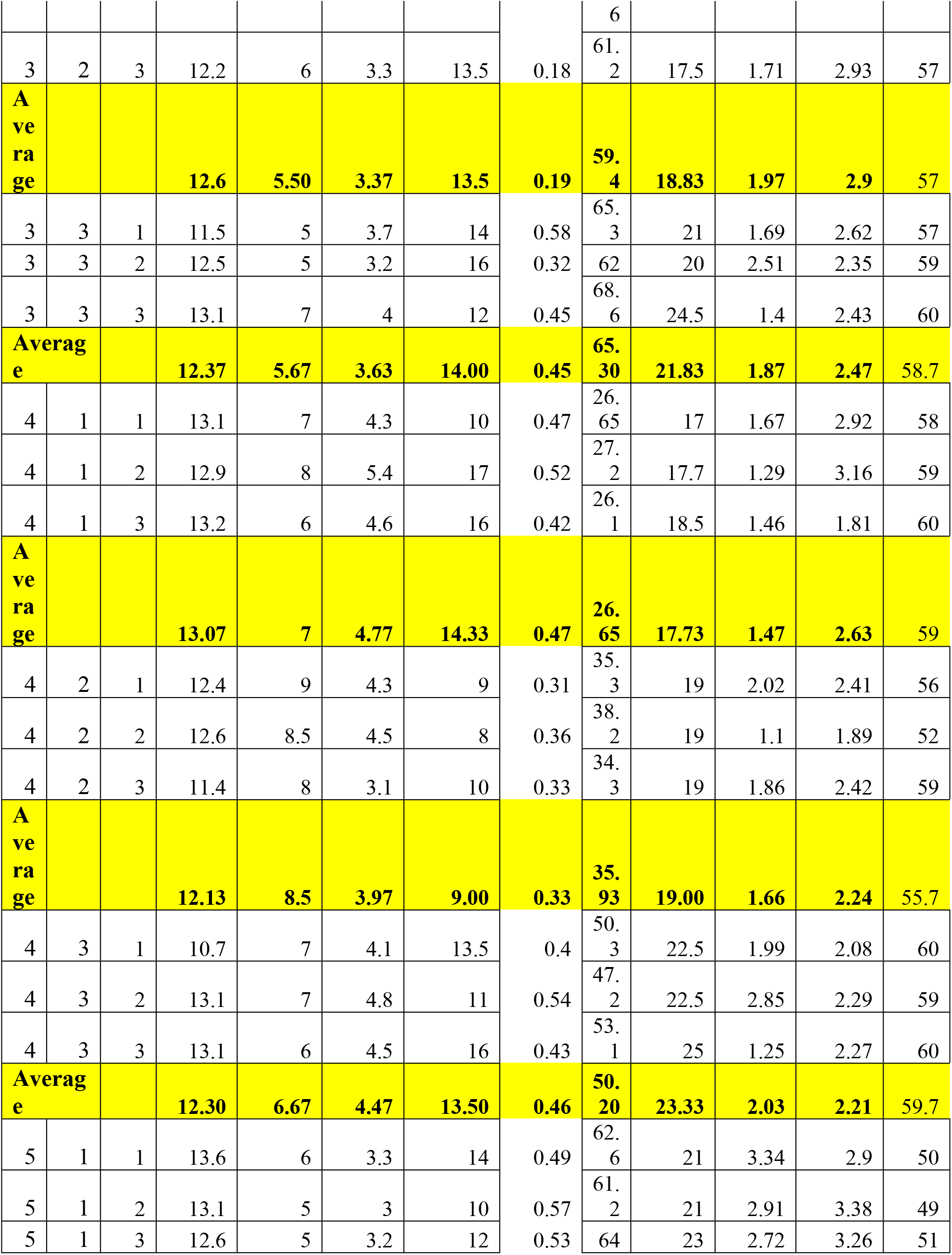
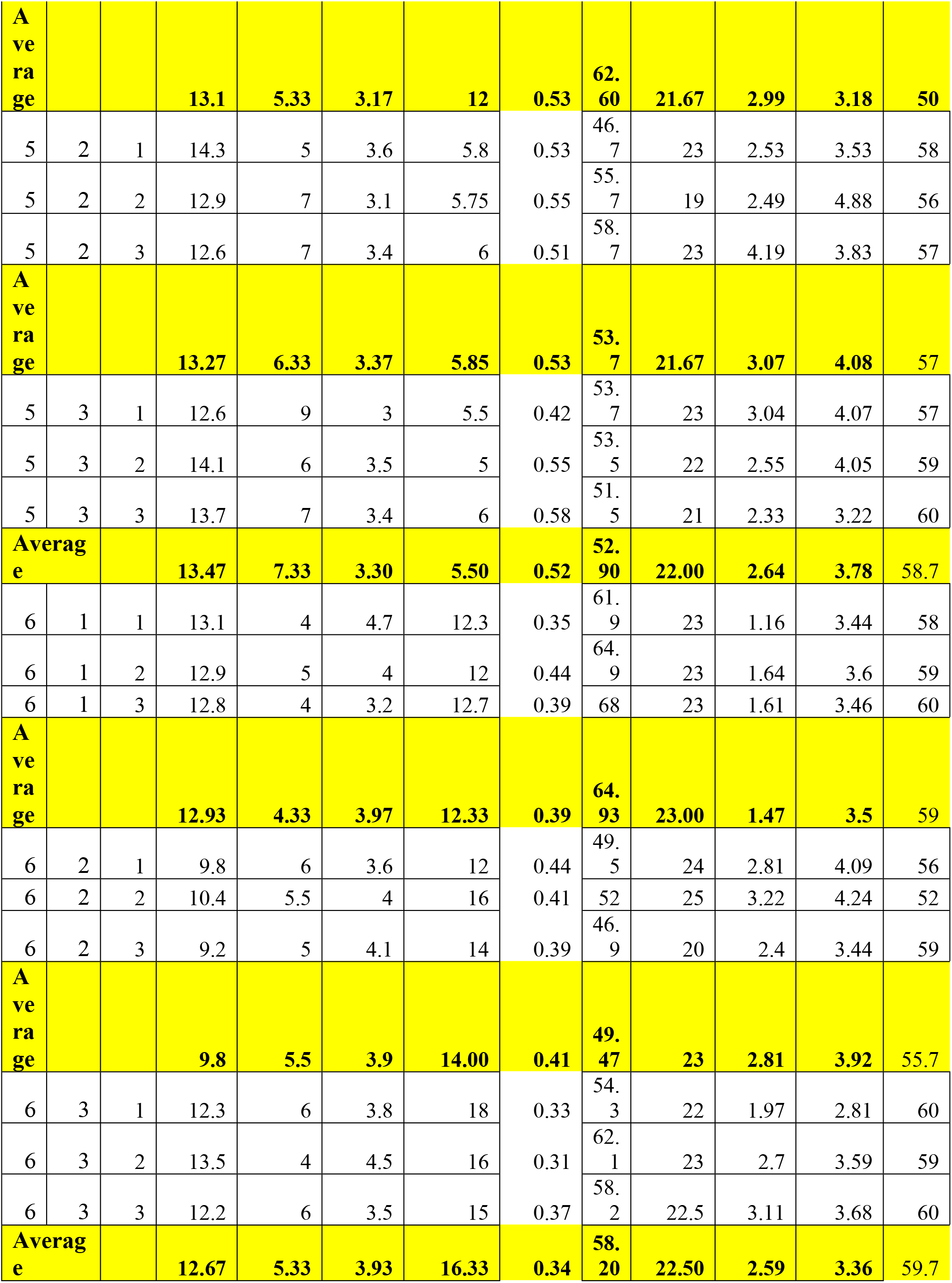
Data of Control.

**Table 4:**
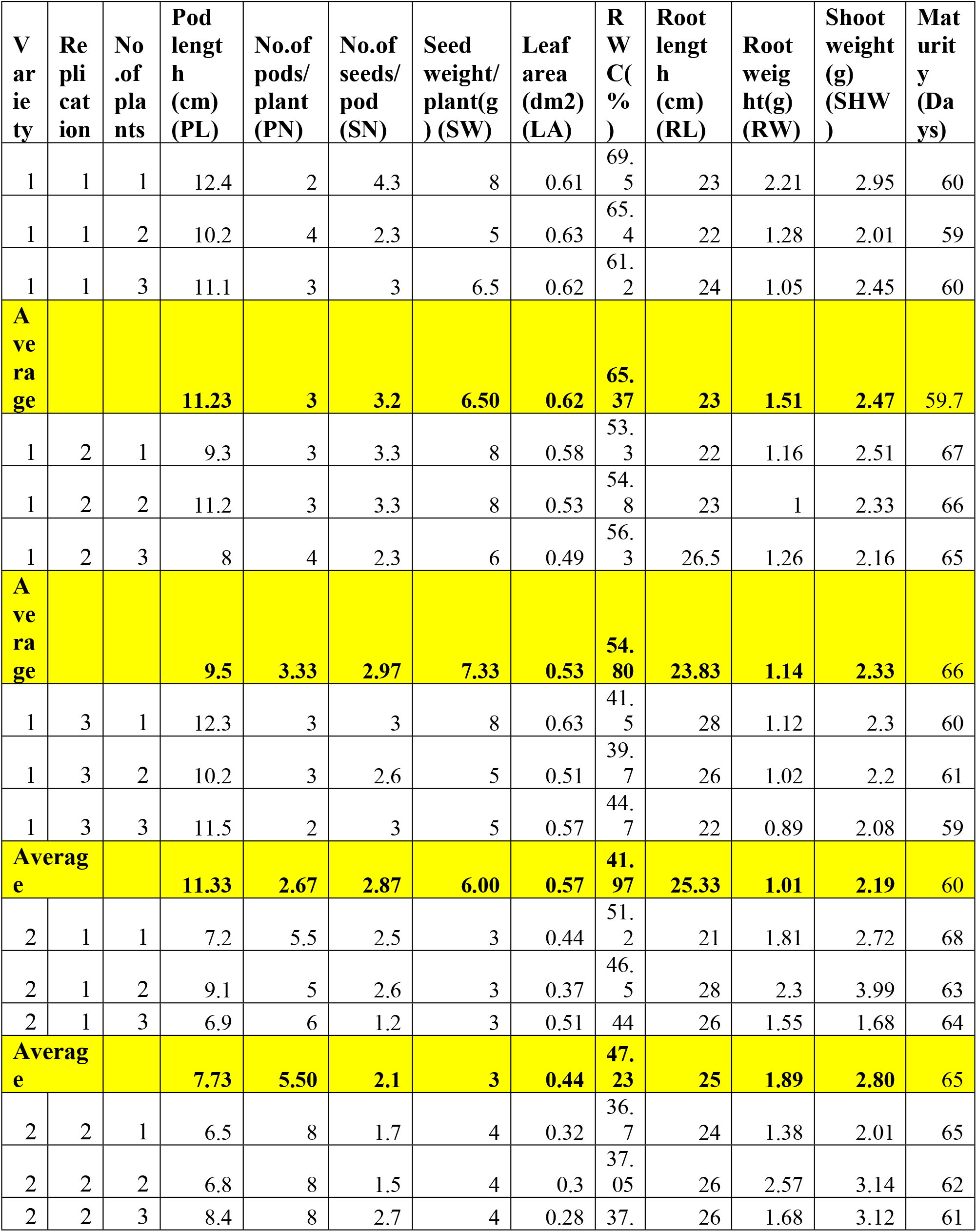
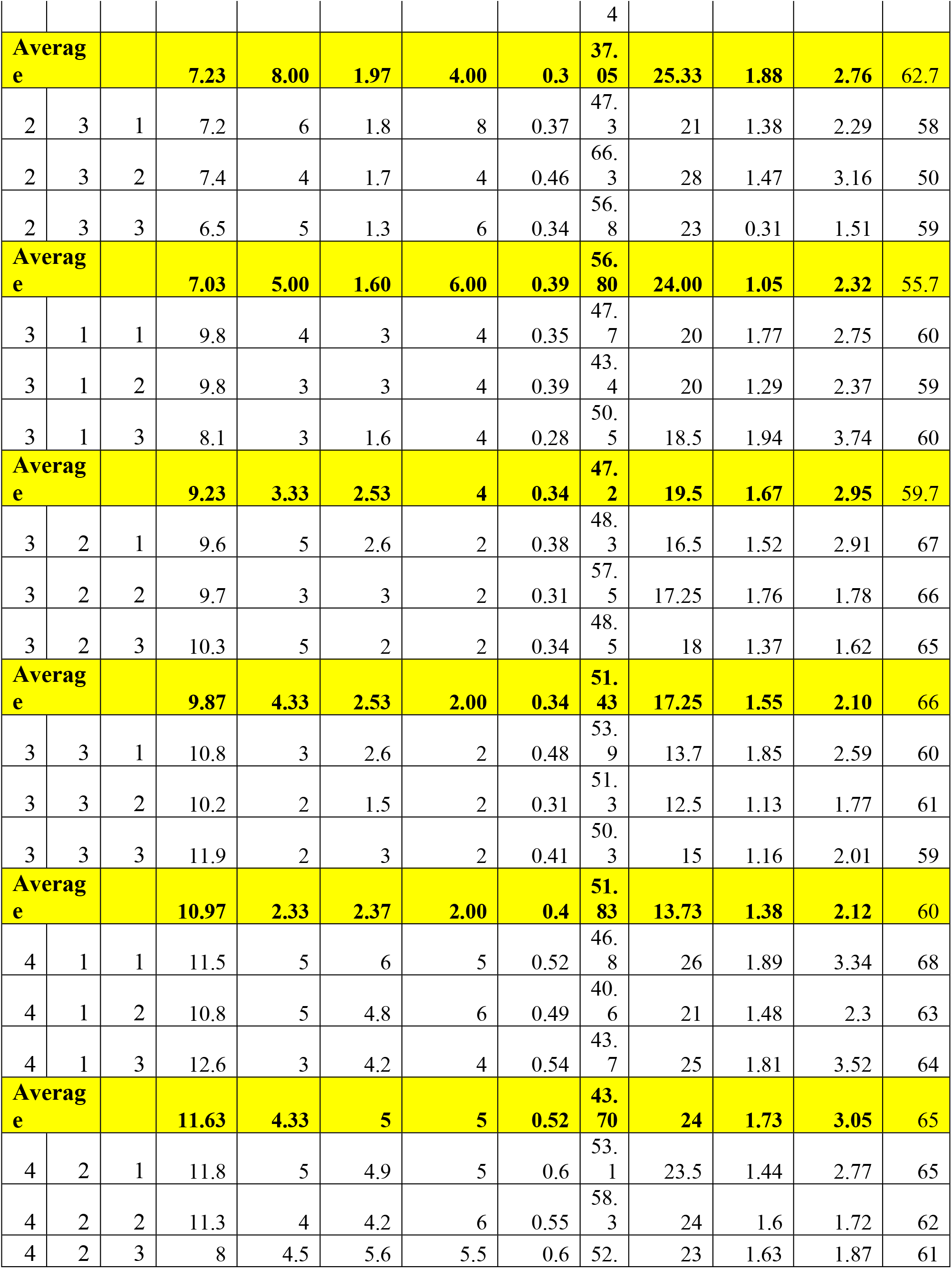
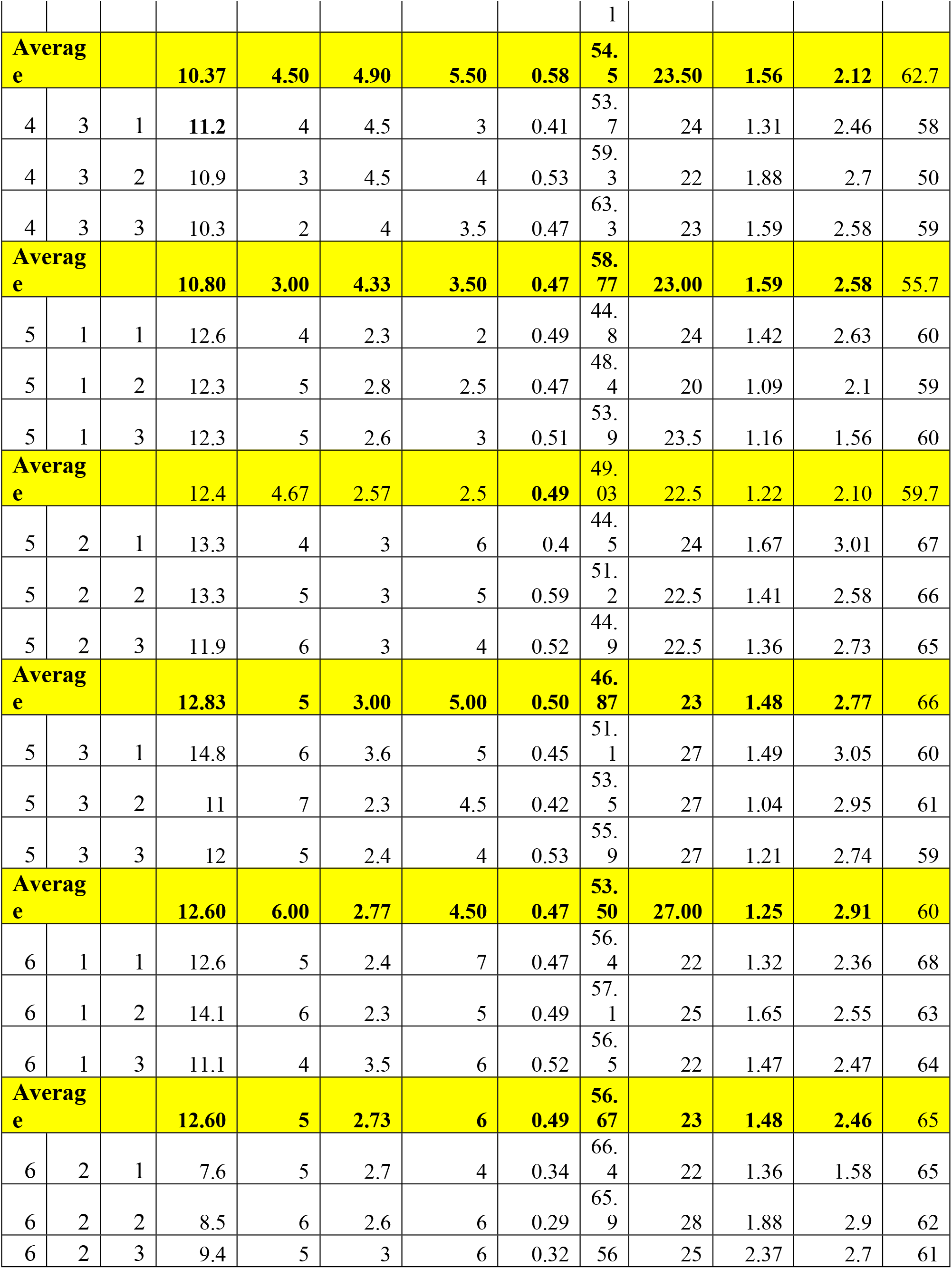
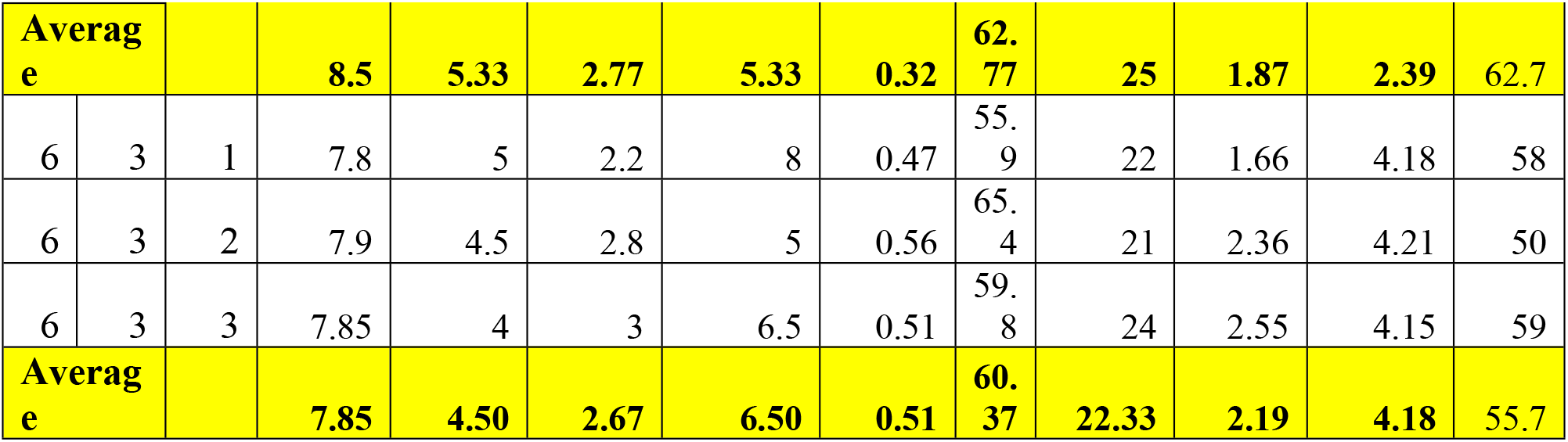
Data of Waterstress.

**Table 5:** Standard Deviation.

| TRT | VAR | REP | PL | PN | SN | SW | LA | RWC | RL | RW | SHW | Maturity |
| --- | --- | --- | --- | --- | --- | --- | --- | --- | --- | --- | --- | --- |
| 1 | 1 | 1 | 12.60 | 3.99 | 4.23 | 12.70 | 0.37 | 46.03 | 28.90 | 2.43 | 2.69 | 50 |
| 1 | 1 | 2 | 12.97 | 4 | 3.17 | 11.00 | 0.79 | 45.86 | 30.00 | 2.16 | 2.68 | 49 |
| 1 | 1 | 3 | 11.70 | 3.99 | 4.63 | 14.50 | 0.50 | 45.7 | 27.80 | 2.29 | 2.67 | 51 |
| SD |  |  | 0.65 | 0.01 | 0.76 | 1.75 | 0.22 | 0.17 | 1.10 | 0.14 | 0.01 | 1.00 |
| 1 | 2 | 1 | 8.60 | 5.33 | 1.63 | 10.27 | 0.49 | 53.97 | 29.14 | 2.41 | 2.30 | 58 |
| 1 | 2 | 2 | 8.24 | 5.67 | 1.93 | 8.5 | 0.43 | 52.30 | 33.00 | 2.69 | 2.32 | 56 |
| 1 | 2 | 3 | 7.88 | 6 | 1.93 | 8.00 | 0.51 | 53.00 | 31.07 | 2.55 | 2.31 | 57 |
| SD |  |  | 0.36 | 0.33 | 0.17 | 1.19 | 0.04 | 0.84 | 1.93 | 0.14 | 0.01 | 1.00 |
| 1 | 3 | 1 | 12.27 | 6.33 | 3.37 | 15.33 | 0.45 | 57.3 | 18.67 | 1.96 | 2.93 | 58 |
| 1 | 3 | 2 | 12.6 | 5.50 | 3.37 | 13.5 | 0.19 | 59.4 | 18.83 | 1.97 | 2.90 | 56 |
| 1 | 3 | 3 | 12.37 | 5.67 | 3.63 | 14 | 0.45 | 65.3 | 21.83 | 1.96 | 2.92 | 57 |
| SD |  |  | 0.17 | 0.44 | 0.15 | 0.95 | 0.15 | 4.15 | 1.78 | 0.01 | 0.02 | 1.00 |
| 1 | 4 | 1 | 13.07 | 7 | 4.77 | 14.33 | 0.47 | 43.06 | 17.73 | 2.02 | 2.63 | 58 |
| 1 | 4 | 2 | 12.13 | 8.5 | 3.97 | 13.90 | 0.33 | 46.63 | 19.00 | 2.01 | 2.53 | 59 |
| 1 | 4 | 3 | 12.3 | 6.67 | 4.47 | 13.50 | 0.46 | 50.2 | 18.00 | 2.03 | 2.58 | 60 |
| SD |  |  | 0.50 | 0.98 | 0.40 | 0.42 | 0.08 | 3.57 | 0.67 | 0.01 | 0.05 | 1 |
| 1 | 5 | 1 | 13.10 | 5.33 | 3.17 | 5.67 | 0.53 | 62.60 | 21.67 | 2.99 | 3.93 | 56 |
| 1 | 5 | 2 | 13.27 | 6.33 | 3.37 | 5.85 | 0.53 | 53.7 | 21.67 | 3.07 | 4.08 | 52 |
| 1 | 5 | 3 | 13.47 | 7.33 | 3.30 | 5.50 | 0.52 | 52.9 | 22.0 | 2.64 | 4.005 | 59 |
| SD |  |  | 0.18 | 1.00 | 0.10 | 0.18 | 0.01 | 5.38 | 0.2 | 0.23 | 0.08 | 3.51 |
| 1 | 6 | 1 | 12.93 | 4.33 | 3.97 | 13.50 | 0.39 | 64.93 | 23.00 | 2.7 | 3.50 | 60 |
| 1 | 6 | 2 | 9.8 | 5.5 | 3.9 | 14.00 | 0.41 | 63.24 | 23 | 2.81 | 3.92 | 59 |
| 1 | 6 | 3 | 12.67 | 5.33 | 3.93 | 16.33 | 0.34 | 61.56 | 22.50 | 2.59 | 3.36 | 60 |
| SD |  |  | 1.74 | 0.63 | 0.03 | 1.51 | 0.04 | 1.69 | 0.29 | 0.11 | 0.29 | 0.58 |
| 2 | 1 | 1 | 11.23 | 3 | 3.2 | 6.50 | 0.62 | 48.38 | 23 | 1.51 | 2.47 | 60 |
| 2 | 1 | 2 | 9.5 | 3.33 | 2.97 | 7.33 | 0.53 | 45.17 | 23.83 | 1.14 | 2.33 | 59 |
| 2 | 1 | 3 | 11.33 | 2.67 | 2.87 | 6.92 | 0.57 | 41.97 | 25.33 | 1.01 | 2.19 | 60 |
| SD |  |  | 1.03 | 0.33 | 0.17 | 0.42 | 0.05 | 3.21 | 1.18 | 0.26 | 0.14 | 0.58 |
| 2 | 2 | 1 | 7.73 | 5.50 | 2.1 | 3 | 0.44 | 39.59 | 25 | 1.89 | 2.80 | 67 |
| 2 | 2 | 2 | 7.23 | 5.25 | 1.97 | 4.00 | 0.30 | 37.05 | 25.33 | 1.88 | 2.76 | 66 |
| 2 | 2 | 3 | 7.03 | 5.00 | 1.6 | 3.5 | 0.39 | 38.32 | 24 | 1.05 | 2.32 | 65 |
| SD |  |  | 0.36 | 0.25 | 0.26 | 0.5 | 0.07 | 1.27 | 0.69 | 0.48 | 0.26 | 1.00 |
| 2 | 3 | 1 | 9.23 | 3.33 | 2.53 | 2 | 0.34 | 47.2 | 19.5 | 1.67 | 2.95 | 60 |
| 2 | 3 | 2 | 9.87 | 4.33 | 2.53 | 2.00 | 0.34 | 48.25 | 17.25 | 1.55 | 2.10 | 61 |
| 2 | 3 | 3 | 10.97 | 2.33 | 2.37 | 2.00 | 0.40 | 49.31 | 18.40 | 1.38 | 2.12 | 59 |
| SD |  |  | 0.88 | 1.00 | 0.10 | 0.00 | 0.03 | 1.06 | 1.13 | 0.14 | 0.49 | 1.00 |
| 2 | 4 | 1 | 11.63 | 4.33 | 5 | 5 | 0.52 | 43.70 | 24 | 1.73 | 3.05 | 68 |
| 2 | 4 | 2 | 10.37 | 4.50 | 4.90 | 5.50 | 0.58 | 46.32 | 23.50 | 1.56 | 2.12 | 63 |
| 2 | 4 | 3 | 10.8 | 3 | 4.33 | 5.25 | 0.47 | 48.95 | 23.00 | 1.59 | 2.58 | 64 |
| SD |  |  | 0.64 | 0.82 | 0.36 | 0.25 | 0.06 | 2.63 | 0.50 | 0.09 | 0.47 | 2.65 |
| 2 | 5 | 1 | 10.8 | 3.00 | 4.33 | 4.75 | 0.49 | 47.69 | 23 | 1.59 | 2.58 | 65 |
| 2 | 5 | 2 | 12.83 | 3.5 | 3.00 | 5.00 | 0.50 | 46.87 | 23 | 1.48 | 2.77 | 62 |
| 2 | 5 | 3 | 12.6 | 4 | 2.77 | 4.50 | 0.47 | 48.52 | 27.00 | 1.25 | 2.91 | 61 |
| SD |  |  | 1.11 | 0.5 | 0.85 | 0.25 | 0.02 | 0.83 | 2.31 | 0.18 | 0.17 | 2.08 |
| 2 | 6 | 1 | 12.60 | 5 | 2.73 | 6 | 0.49 | 56.67 | 23 | 1.48 | 2.46 | 58 |
| 2 | 6 | 2 | 8.5 | 5.33 | 2.77 | 5.33 | 0.32 | 58.52 | 25 | 1.87 | 2.39 | 50 |
| 2 | 6 | 3 | 7.85 | 4.50 | 2.67 | 6.5 | 0.51 | 57.59 | 22.33 | 2.19 | 2.41 | 59 |
| SD |  |  | 2.58 | 0.42 | 0.05 | 0.59 | 0.10 | 0.93 | 1.39 | 0.36 | 0.03 | 4.93 |
| TRT 1= Control; TRT 2 = Stress |  |  |  |  |  |  |  |  |  |  |  |  |

**Table 6:**
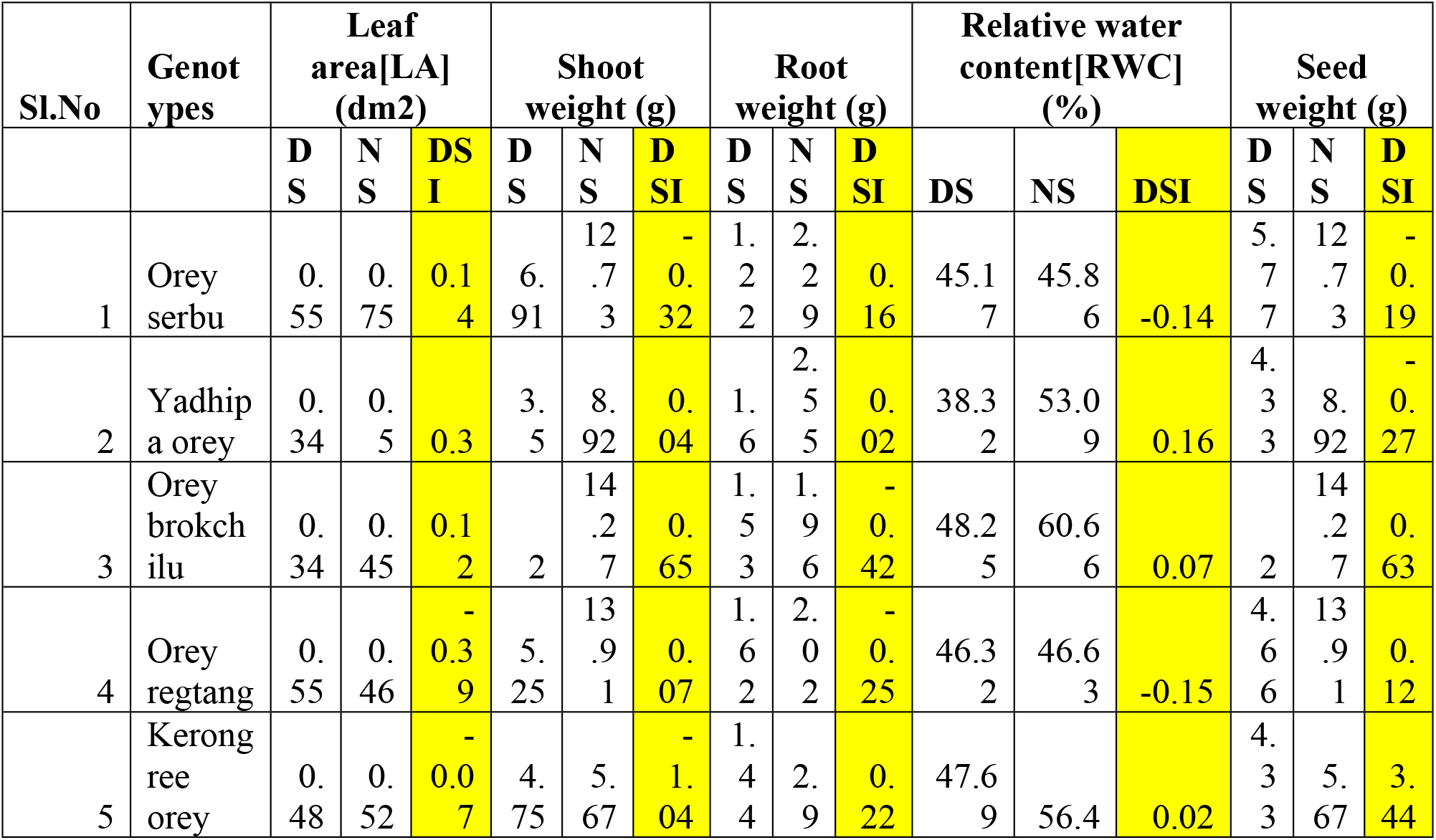
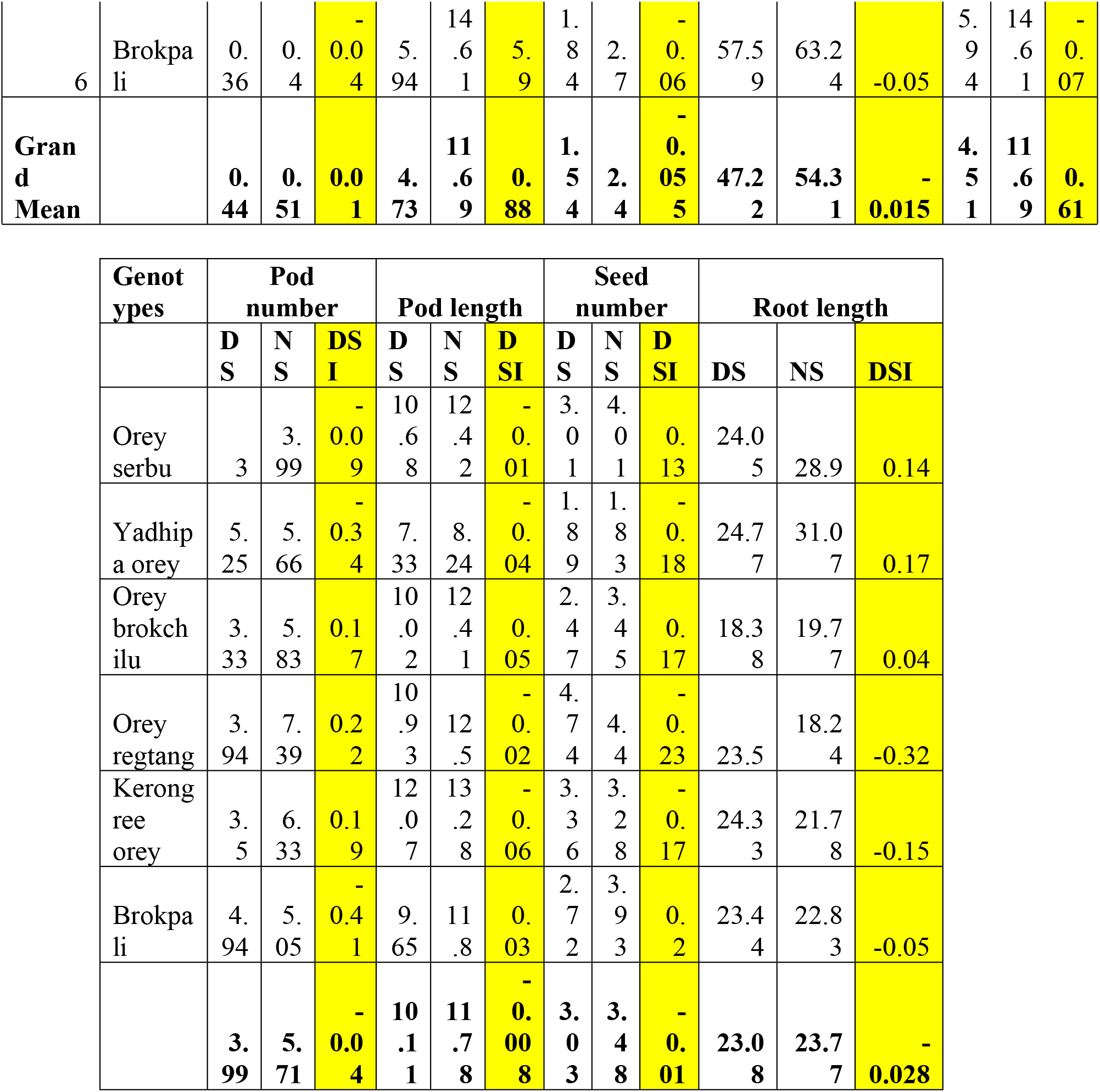
DSI Data.

**Table 7:**
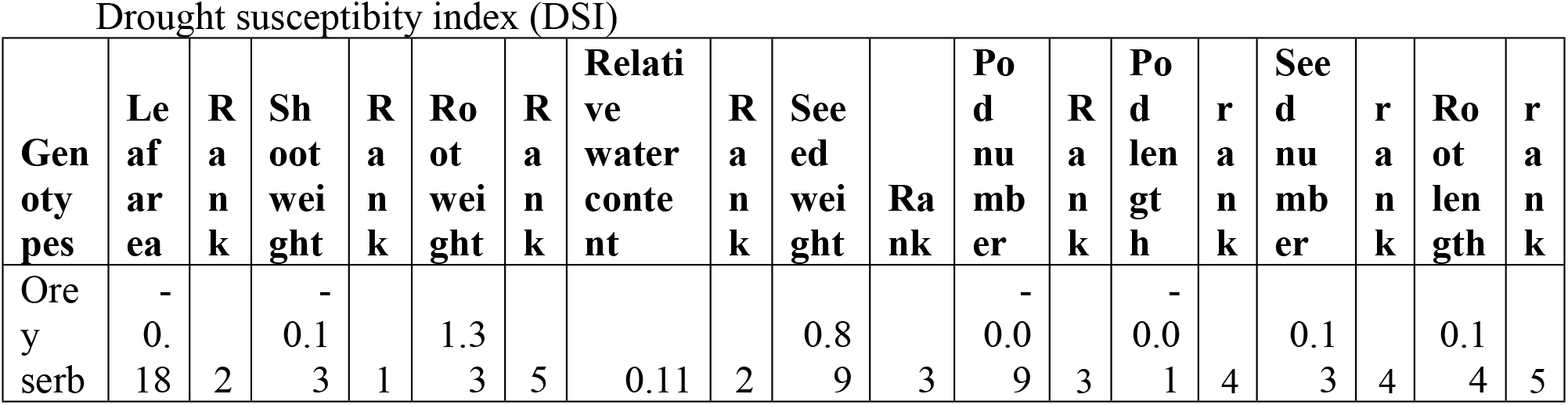
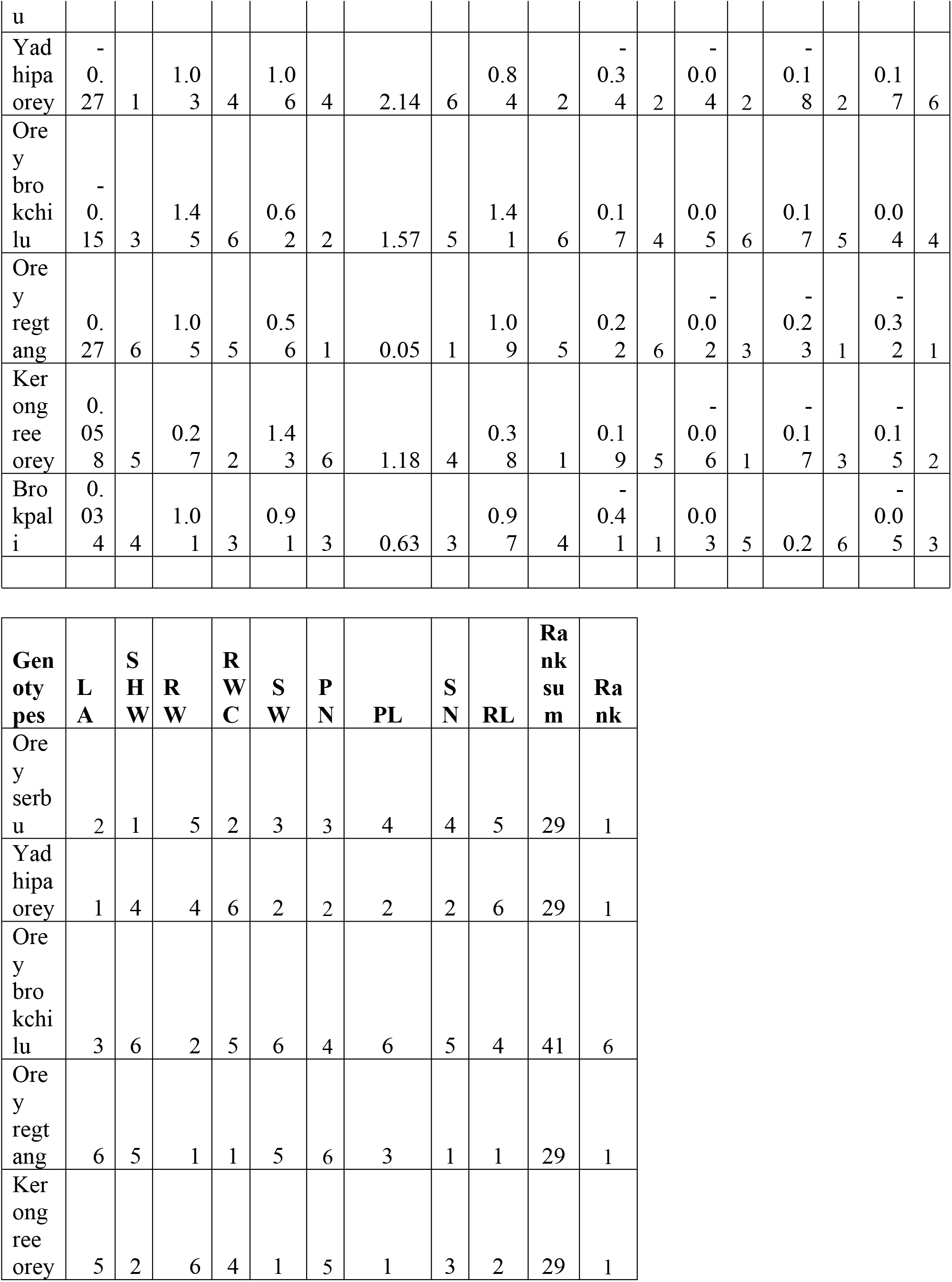
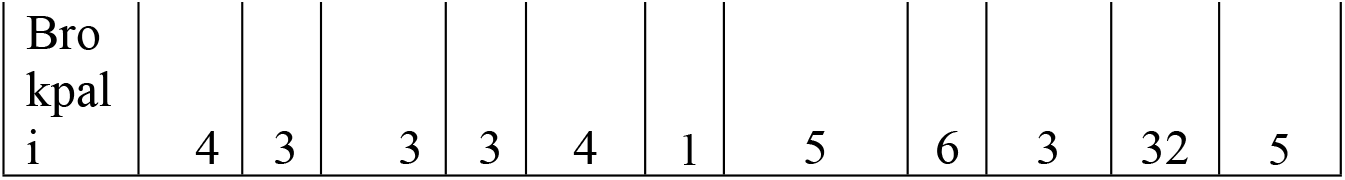
Rank of DSI.

**Table 8:** Data used for genereation graphs.

| Leaf area (dm<sup>2</sup>) |  |  |
| --- | --- | --- |
| Genotypes | Non stress | Drought stress |
| Orey serbu | 0.75 | 0.55 |
| Yadhipa orey | 0.5 | 0.34 |
| Orey brokchilu | 0.45 | 0.34 |
| Orey regtang | 0.46 | 0.38 |
| Keronngree orey | 0.52 | 0.4 |
| Brokpali | 0.4 | 0.3 |
| **Mean** | **0.51** | **0.43** |
| **SD** | **0.12** | **0.09** |
| **CV%** | **24.16** | **20.47** |
| **SE** | **0.05** | **0.04** |

| Pod length (cm) |  |  |
| --- | --- | --- |
| Genotypes | Non stress | Drought stress |
| Orey serbu | 12.42 | 10.68 |
| Yadhipa orey | 8.24 | 7.33 |
| Orey brokchilu | 12.41 | 10.02 |
| Orey regtang | 12.5 | 10.93 |
| Keronngree orey | 13.28 | 12.07 |
| Brokpali | 11.8 | 9.65 |
| **Mean** | **11.78** | **10.11** |
| **SD** | **1.79** | **1.60** |
| **CV%** | **15.24** | **15.81** |
| **SE** | **0.73** | **0.65** |

Pod number
| **Genotypes** | **Non stress** | **Drought stress** |
| --- | --- | --- |
| Orey serbu | 3.99 | 3 |
| Yadhipa orey | 5.66 | 5.25 |
| Orey brokchilu | 5.83 | 3.33 |
| Orey regtang | 7.39 | 3.94 |
| Keronngree orey | 6.33 | 3.5 |
| Brokpali | 5.05 | 4.94 |
| **Mean** | **5.71** | **3.99** |
| **SD** | **1.15** | **0.91** |
| **CV%** | **20.15** | **22.83** |
| **SE** | **0.47** | **0.37** |

Root length (cm)
| **Genotypes** | **Non stress** | **Drought stress** |
| --- | --- | --- |
| Orey serbu | 28.9 | 24.05 |
| Yadhipa orey | 31.07 | 24.77 |
| Orey brokchilu | 19.77 | 18.38 |
| Orey regtang | 18.24 | 23.5 |
| Keronngree orey | 21.78 | 24.33 |
| Brokpali | 22.83 | 23.44 |
| **Mean** | **23.77** | **23.08** |
| **SD** | **5.12** | **2.36** |
| **CV%** | **21.54** | **10.21** |
| **SE** | **2.09** | **0.96** |

Root weight (g)
| **Genotypes** | **Non stress** | **Drought stress** |
| --- | --- | --- |
| Orey serbu | 2.29 | 1.22 |
| Yadhipa orey | 2.55 | 1.60 |
| Orey brokchilu | 1.96 | 1.53 |
| Orey regtang | 2.02 | 1.62 |
| Keronngree orey | 2.9 | 1.44 |
| Brokpali | 2.7 | 1.84 |
| **Mean** | **2.40** | **1.54** |
| **SD** | **0.38** | **0.21** |
| **CV%** | **15.73** | **13.39** |
| **SE** | **0.15** | **0.08** |

RWC (%)
| **Genotypes** | **Non stress** | **Drought stress** |
| --- | --- | --- |
| Orey serbu | 45.86 | 45.17 |
| Yadhipa orey | 53.09 | 38.32 |
| Orey brokchilu | 60.66 | 48.25 |
| Orey regtang | 46.63 | 46.32 |
| Keronngree orey | 56.4 | 47.69 |
| Brokpali | 63.24 | 57.59 |
| **Mean** | **54.31** | **47.22** |
| **SD** | **7.16** | **6.21** |
| **CV%** | **13.18** | **13.16** |
| **SE** | **2.92** | **2.54** |

Shoot weight (g)
| **Genotypes** | **Non stress** | **Drought stress** |
| --- | --- | --- |
| Orey serbu | 12.73 | 6.91 |
| Yadhipa orey | 8.92 | 3.50 |
| Orey brokchilu | 14.27 | 2 |
| Orey regtang | 13.91 | 5.25 |
| Keronngree orey | 5.67 | 4.75 |
| Brokpali | 14.61 | 5.94 |
| **Mean** | **11.68** | **4.71** |
| **SD** | **3.61** | **1.76** |
| **CV%** | **30.89** | **37.32** |
| **SE** | **1.47** | **0.72** |

Seed number
| **Genotypes** | **Non stress** | **Drought stress** |
| --- | --- | --- |
| Orey serbu | 4.01 | 3.01 |
| Yadhipa orey | 1.83 | 1.89 |
| Orey brokchilu | 3.45 | 2.47 |
| Orey regtang | 4.4 | 4.74 |
| Keronngree orey | 3.28 | 3.36 |
| Brokpali | 3.93 | 2.72 |
| **Mean** | **3.48** | **3.03** |
| **SD** | **0.90** | **0.97** |
| **CV%** | **26.00** | **32.14** |
| **SE** | **0.37** | **0.40** |

Seed weight (g)
| **Genotypes** | **Non stress** | **Drought stress** |
| --- | --- | --- |
| Orey serbu | 12.73 | 5.77 |
| Yadhipa orey | 8.92 | 4.33 |
| Orey brokchilu | 14.27 | 2 |
| Orey regtang | 13.91 | 4.66 |
| Keronngree orey | 5.67 | 4.33 |
| Brokpali | 14.61 | 5.94 |
| **Mean** | **11.68** | **4.50** |
| **SD** | **3.61** | **1.42** |
| **CV%** | **30.89** | **31.46** |
| **SE** | **1.47** | **0.58** |

Maturity
| **Genotypes** | **Non stress** | **Drought stress** |
| --- | --- | --- |
| Orey serbu | 50 | 60 |
| Yadhipa orey | 57 | 66.00 |
| Orey brokchilu | 58.6 | 59.6 |
| Orey regtang | 59 | 65 |
| Keronngree orey | 55.6 | 62 |
| Brokpali | 59.6 | 55.6 |
| **Mean** | **56.60** | **61.50** |
| **SD** | **3.56** | **4.22** |
| **CV%** | **6.29** | **6.85** |
| **SE** | **1.45** | **1.89** |

